# Influence of anatomical features of different brain regions on the spatial localization of fiber photometry signals

**DOI:** 10.1101/2021.08.02.454755

**Authors:** Cinzia Montinaro, Marco Pisanello, Marco Bianco, Barbara Spagnolo, Filippo Pisano, Antonio Balena, Francesco De Nuccio, Dario Domenico Lofrumento, Tiziano Verri, Massimo De Vittorio, Ferruccio Pisanello

## Abstract

Fiber photometry is widely used in neuroscience labs for *in vivo* detection of functional fluorescence from optical indicators of neuronal activity with a simple optical fiber. The fiber is commonly placed next to the region of interest to both excite and collect the fluorescence signal. However, the path of both excitation and fluorescence photons is altered by the uneven optical properties of the brain, due to local variation of the refractive index, different cellular types, densities and shapes. Nonetheless, the effect of the local anatomy on the actual shape and extent of the volume of tissue that interfaces with the fiber has received little attention so far. To fill this gap, we measured the size and shape of fiber photometry efficiency field in the primary motor and somatosensory cortex, in the hippocampus and in the striatum of the mouse brain, highlighting how their substructures determine the detected signal and the depth at which photons can be mined. Importantly, we show that the information on the spatial expression of the fluorescent probes alone is not sufficient to account for the contribution of local subregions to the overall collected signal, and it must be combined with the optical properties of the tissue adjacent to the fiber tip.

## INTRODUCTION

The development of high-efficiency optical indicators of neural activity has widened the application of fiber photometry (FP) [1–4], a method employing flat-cleaved step-index optical fibers (OFs) to monitor time-dependent functional fluorescence and/or lifetime variations related to several physiological phenomena, including calcium (Ca^2+^) levels [5], membrane potential [6], neurotransmitters transients [7] and the intracellular biochemical state of neurons [8]. The OF is commonly placed next to the region of interest and used to excite the fluorescent indicators and to collect the resulting functional signal. The brain volume contributing to the overall signal depends on the constitutive parameters of the OF, including numerical aperture (NA), core/cladding dimensions and refractive index [9], and it is the result of the combination of the three-dimensional excitation and collection fields [9–12]. Moreover, both excitation and fluorescence photons undergo tissue attenuation and scattering, and the generated fluorescence strongly depends on: (i) how the excitation light distributes at the output of the OF, and (ii) how many fluorescence photons generated in a specific point reach the fiber facet.

While different methods exist to estimate the FP sensitivity volume in brain tissue [9,13], the optical properties of the brain are highly uneven, not only at the cellular and subcellular level, but also on the scale of hundreds of micrometers and millimeters. The anatomical distribution of cells bodies, for instance, significantly varies across different brain regions, and distinct structures are characterized by a different cell density, while others contain mostly neuropil. In this regard, representative examples can be identified in the cerebral cortex (CTX), the hippocampus (HP) and the striatum (STR) of the mouse brain. The CTX has a columnar structure consisting of six alternating layers (LI-LVI) [14], each one with a specific anatomy characterized by different cellular densities and cell types. In addition, the depth of each layer and its composition depends on the specific subregion, with peculiar known differences across motor, somatosensory, associative and visual cortex [15]. Similarly, the HP has a layered structure too, with neural bodies mainly concentrated along a curve, from *Cornu Ammonis* 1 (CA1) through *Cornu Ammonis* 4 (CA4) [16,17], with basal and apical dendrites extending in two different directions and generating highly fibrous regions above and below the cell bodies layers. The STR organization follows, instead, a spatio-molecular code and the striatal circuitry can be divided into two major pathways of striatal projection neurons (SPNs) that have distinct neuroanatomical and molecular features [18,19]. Therefore, the anatomy of cellular shapes and their distribution adds an additional layer of complexity to the problem of estimating the spatial localization of FP signal. At the same time, the use of genetically encoded fluorescent indicators of neural activity makes a subpopulation of cells act as source of functional fluorescence, while non-tagged neurons influence the collected signal as a passive optical medium, defining the optical properties of the tissue.

In this study we report how the shape and size of the three-dimensional fiber photometry efficiency field changes across the cortex, hippocampus and striatum of the mouse brain. We used the widely adopted *Thy1*-GCaMP6 line [20,21]. This strain is characterized by GCaMP expression under the *Thy1* promoter, an immunoglobulin superfamily expressed by projection neurons, allowing for identifying neuronal somata and projections across different brain areas [22–24]. Importantly, we relate the measured FP efficiency fields to the local cytoarchitecture specific of the investigated regions, highlighting significant intra-region differences. In this framework, our results suggest that fiber photometry data should be analyzed by considering the specific region from which the collected signal is generated, since each peculiar substructure contributes in defining the final sensitivity volume in terms of both size and shape.

## RESULTS

### Optical setup and methodology

A two-photon (2P) laser scanning system, displayed in *Figure 1A*, was used to measure the illumination (*β*) and collection (*η*) fields [9] of a OF stub with NA = 0.39 and core diameter of 200 μm, positioned next to the region of interest on 300 μm-thick coronal brain slices obtained from *Thy1*-GCaMP6*s* transgenic mice (see *right* inset *Figure 1A*). A fs-pulsed near-infrared (NIR) laser beam (λ_ex_ = 920 nm) was used to generate a fluorescent voxel scanned in three dimensions close to the facet of the fiber, in a z-stack spanning 100 µm across the fiber facet (z step 10 µm). Resulting fluorescence was detected by a photomultiplier tube (“*μscope* PMT”, *μ*) in non-descanned epifluorescence configuration, and simultaneously a fraction of the signal was collected by the optical fiber and guided to a second PMT (“*fiber* PMT”, *f*). This generated two images stacks *μ(x,y,z)* and *f(x,y,z)*, respectively. The OF’s collection efficiency field was then computed as *η(x,y,z)*= *f(x,y,z)/μ(x,y,z)* [9]. The same system was employed to measure the normalized excitation field *β*(*x,y*) in the same brain region by delivering a 473 nm continuous wave (CW) laser beam through the same fiber and collecting the resulting fluorescence signal with a sCMOS camera. Since the depth of focus of the employed objective (Olympus XLFLUOR-340 4x/NA 0.28) was estimated to be 57 µm [25], *β* was acquired as a single slice.

**Figure 1:**
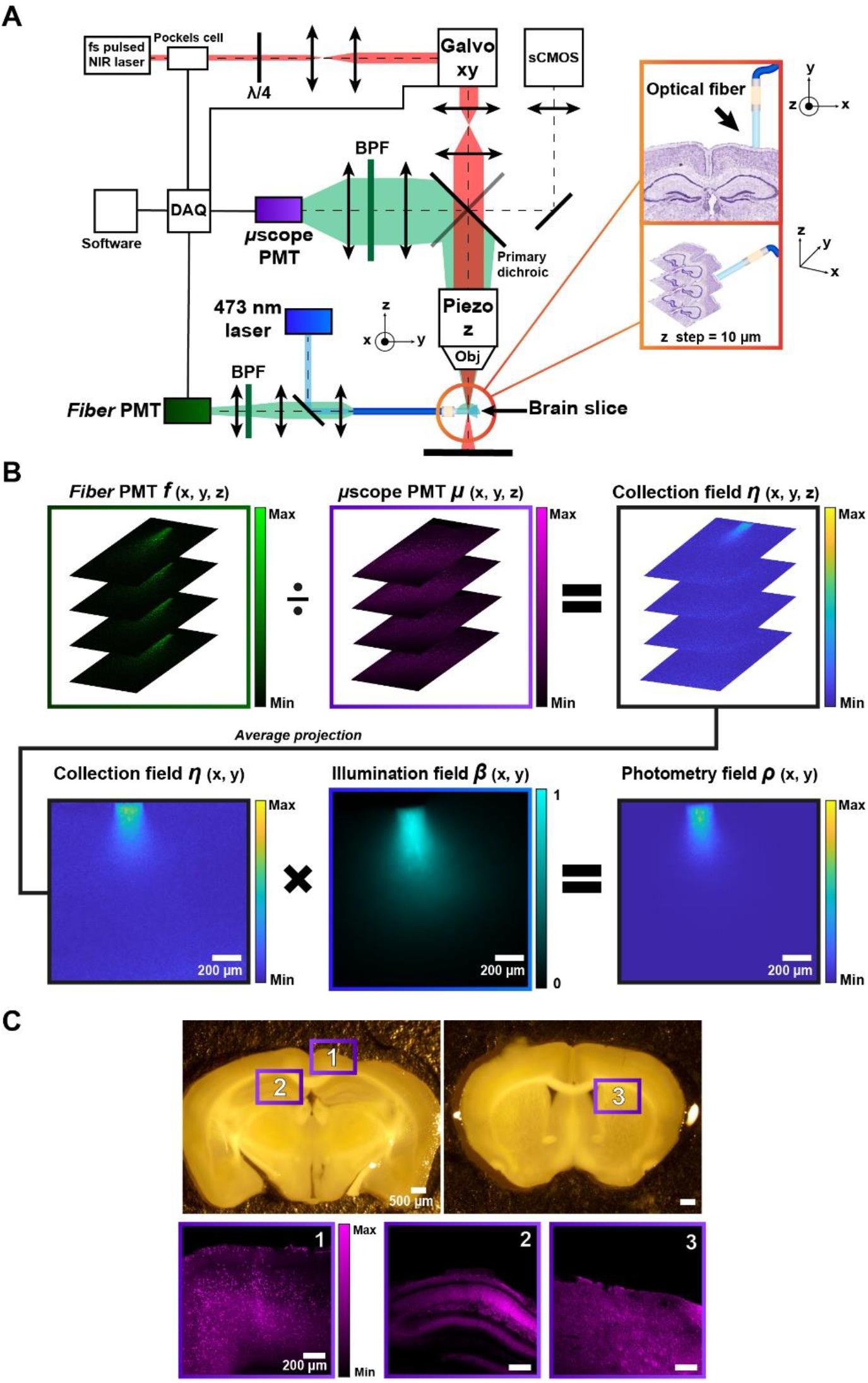
(**A**) Setup used to measure the collection and illumination fields of an optical fiber placed next to the brain region of interest. (**B**) Schematic representation of the combination of *η(x,y) and β(x,y)* to obtain *ρ(x,y)*. (**C**) *(top)* Stereomicroscope images of coronal brain slices obtained from *Thy1*-GCaMP6*s* transgenic mice, *(bottom)* two-photon microscope images of the brain regions: (1) CTX, (2) HP, (3) STR. The boxes’ colors (magenta, green and blue) in (**B**) and in (**C**) correspond to the detectors, shown in (**A**).

To match *η* to *β*, the average projection on six slices of *η(x,y,z)* (equivalent to a thickness of 60 µm) was performed, resulting in the 2D field *η(x,y)*. The photometry efficiency field was then retrieved as *ρ(x,y)*= *η(x,y)β(x,y)* [9,13]. This overall procedure is summarized in *Figure 1B. ρ(x,y)* therefore takes into account the optical fiber’s constitutive parameters and the properties of the brain tissue (i.e. refractive index, absorption and scattering coefficients) interposed between the light source and the OF [9,26]. We employed *ρ* as main figure of merit to evaluate the dependence of the FP signal in the different regions investigated in this work. *Figure 1C (top)* shows representative coronal brain slices obtained from *Thy1*-GCaMP6*s* transgenic mice highlighting the specific brain regions investigated in this work, together with their two-photon microscope images *(bottom)*.

### Comparing photometry efficiency fields across CTX, HP and STR

To identify the influence of anatomical features on the spatial behavior of FP signals, we have chosen to investigate light collection volumes of optical fibers placed next to the mouse cerebral cortex, the hippocampus and the striatum, which are characterized by intrinsically different cytoarchitecture.

The cerebral cortex can be divided in different areas: motor, somatosensory, visual and auditory, each of which has its own function and organization. All the neocortical areas derived from a six-layered structure where the term layer refers to an aggregate of neuronal cell bodies and/or neuropil. As a particular case, primary motor cortex (M1) lacks LIV, consisting of granular cells [14]. Moreover, the cerebral cortex shrinks into a single S-shaped layer to constitute the hippocampus, in which, however, the subdivision into substructures continues, distinguishing CA1, CA2, CA3 and CA4 areas [17]. The main output connections of the hippocampus are represented by the pyramidal neurons of the CA1, innervating numerous areas of the brain, with a non-homogeneous cell typology. CA1 pyramidal neurons can be differentiated according to their size, shape and location of their soma, or to basal and apical dendritic arborizations and specializations [27]. On the contrary, striatum cellular organization appears to be more homogeneous, but it presents a peculiar distribution into patch and matrix compartments and in two main pathways, formed by subtypes of striatal projection neurons (SPN) having different molecular identities, and featuring a subdivision based on a spatio-molecular code [19].

*Figure 2A* shows representative results of *f* and *β* measured for a 0.39 NA OF placed next to M1, CA1 and dorsal STR, together with a reference 2P fluorescence image. In the case of M1 (*Figure 2A, left column*), both collection and excitation fields extend up to LV, with some signal arising also from LVI, with decreasing intensity as a function of depth. However, the photometry efficiency field *ρ* has shorter extension in depth with respect to *η* and *β*, being the product between the two. This is shown in *Figure 2B, top*, showing iso-intensity lines at 10%, 20%, 40%, 60% and 80% of the maximum number of collected photons on the photometry field, together with their 3D representation in a rotationally symmetric diagram (*Figure 2B, bottom*). Isolines in *Figure 2B top* have a narrow and elongated shapes with: isolines at 80% and 60% extending at the boundary between LI and LII-III, isolines at 40% that do not exceed the depth of LII-III, and isolines at 20% and 10% that reach up to LV. In HP (*Figure 2A-B, central column*), the isolines clearly follow the anatomical structure of CA1 with a two-lobe shape: isolines from 80% to 40% outline the basal dendrites, while 20% and 10% isolines reach the apical dendrites layer and slightly narrow across the cell bodies of pyramidal neurons. In STR, instead, no clear anatomy-dependent collection is observed, while the spatial behavior of the photometry efficiency field results to be distributed more homogeneously.

**Figure 2:**
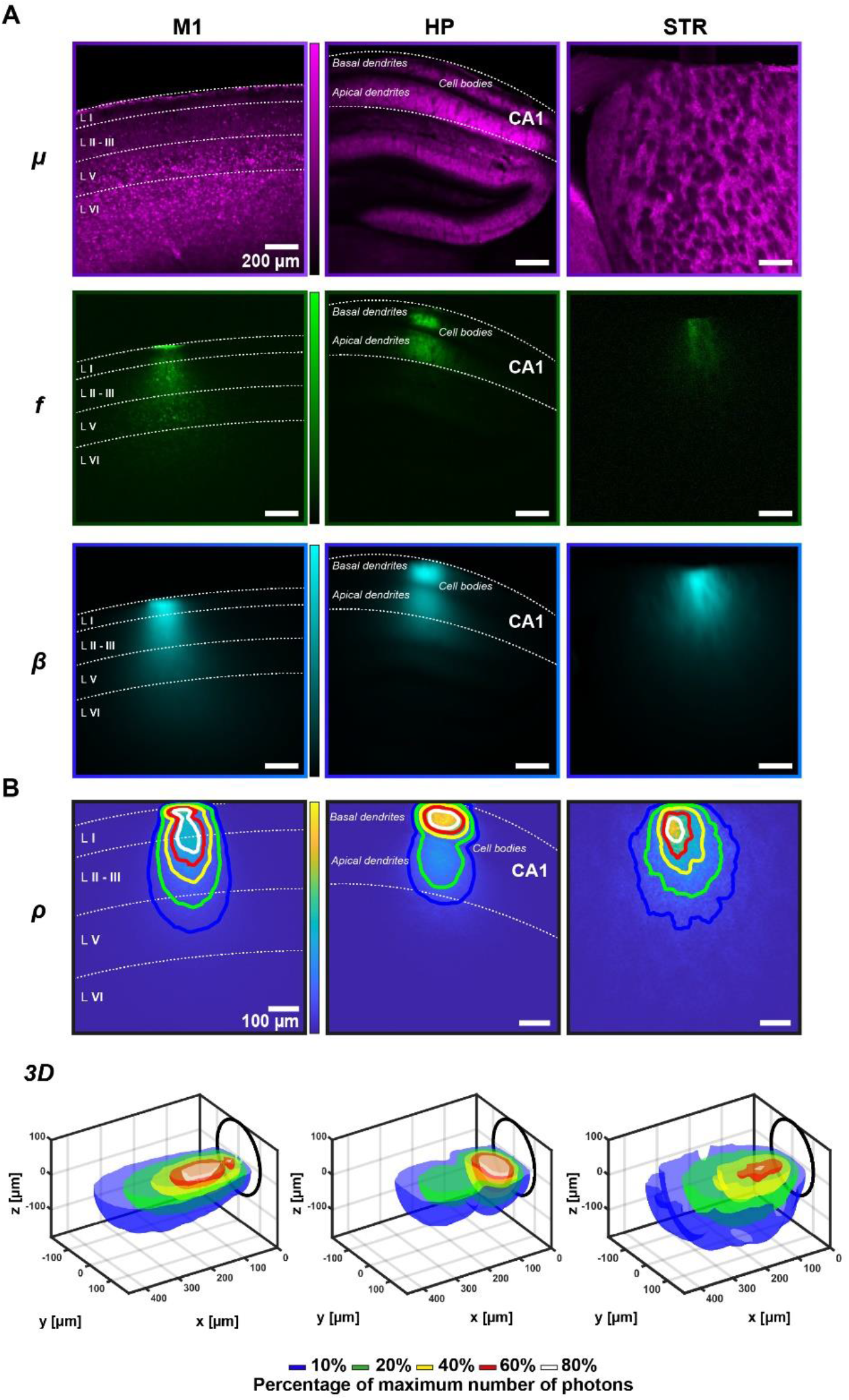
(**A**) Representative *µscope* PMT image (*top*), *fiber* PMT image (*middle*) and the illumination field (*bottom*) of primary motor cerebral cortex (*left*), hippocampus (*center*) and striatum (*right*). Scalebar in all the panels (**A**) is 200 µm. (**B**) *(top)* Photometry collection efficiency field for each region with comparison of iso-intensity surfaces at 10%, 20%, 40%, 60%, and 80% of the maximum number of photons are shown (in blue, green, yellow, red, white respectively); *(bottom)* their 3D configuration as surfaces of revolution obtained by rotating the isolines around the fiber axis. Scalebar in all the panels (**B**) (*top*) is 100 µm. Images of *µ* and *f* in panel (**A**) were adjusted for visualization sake.

The difference between the detection depths at the different contribution percentages is quantified in *Figure 3A*: each horizontal line represents the maximum depth 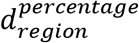 reached by a specific iso-intensity surface. The blue data points, relative to, 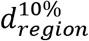, clearly show that the photometry efficiency field extends deeper for M1 and STR with respect to HP (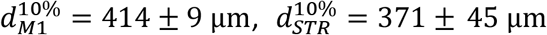 and 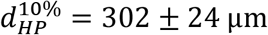, mean ± standard deviation on n = 3, single measures reported in *Supplementary Figures 1-3*).

**Figure 3:**
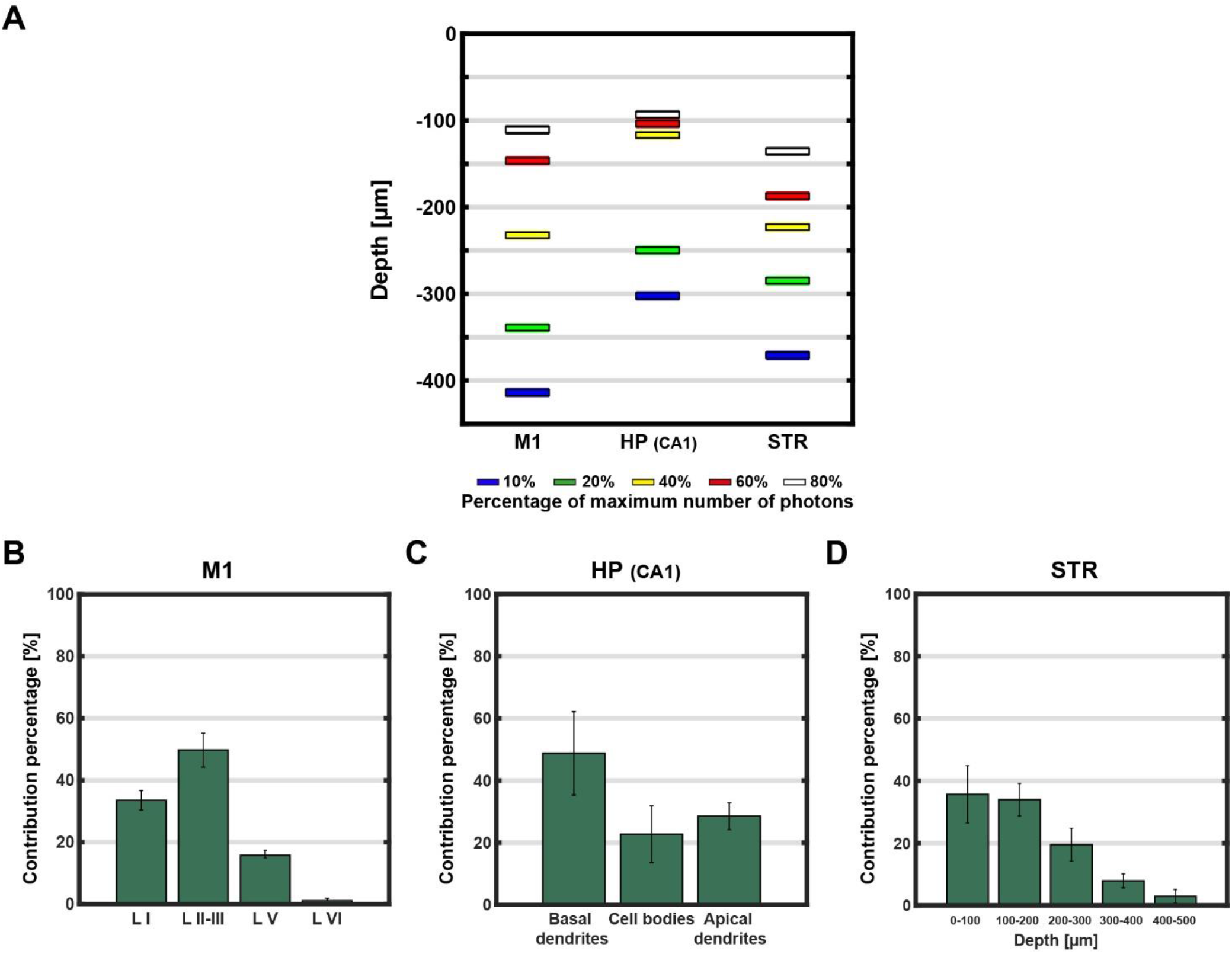
(**A**) Detection depths at the different contribution percentages between M1, HP and STR. Each bar indicates the different contribution of each anatomical feature in (**B**) M1, (**C**) CA1 subcellular organization reveals three different areas contributing to the overall photometry signal: basal dendrites, cell bodies and apical dendrites and (**D**) STR. Error bars in bar graphs in panels (**B**), (**C**) and (**D**) represent the standard deviation of the percentages evaluated on N = 3 brain slices.

Another peculiar difference is that in the HP the high intensity region (see white, red and yellow lines) is confined above a depth of ∼117 µm, while for M1 and STR it is more evenly distributed until ∼232 µm and ∼223 µm respectively (see the spacing between red, yellow and white data points in *Figure 3A*).

On these bases, we have estimated how much each anatomical and cellular feature contributes to the collected signal. This is reported in *Figure 3B* and *C* for M1 and HP. In the case of M1 roughly half of the signal is recorded from LII-III, despite GCaMP is mostly expressed by LV pyramidal neurons [28]. Therefore, when the fiber is placed at the cortex surface, the collected fluorescence is mostly generated by the apical dendrites extending from LV’s soma to LII-III and LI, which together account for ∼80% of the overall collected photons against the ∼16% assigned to LV. In HP instead (*Figure 3C*) the fluorescence signal generated in CA1 has a peculiar sub-distribution: ∼49% derives from the top layer constituted by basal dendrites, ∼23% from pyramidal cell bodies layer and the remaining ∼28% can be ascribed to the apical dendrites region. The striatum is instead macroscopically more uniform, and its microscopic anatomy is concealed by histochemical cells organization in striosomes (or patches) and matrix compartments. Indeed, STR has a non-layered cytoarchitecture and in addition it receives several afferent fibers from cortical and subcortical structure and projects efferent fibers to basal ganglia nuclei [29]. This results in a homogeneous propagation of photons, and the spatial distribution of photometry efficiency can hardly be related to specific anatomical features. Instead, it is interesting to analyze it as a function of depth, as clearly highlighted in the progressive signal decrease in the bar graph in *Figure 3D*.

### Variability of photometry efficiency field across motor and somatosensory cortex

The measurements described in Section 2.2 suggest that the different anatomy of functional brain structures influences the shape of the photometry efficiency field, as well as the maximum depth at which the signal is collected; therefore, we expect that also anatomical differences within the same region strongly affect the spatial behavior of *ρ*. A representative example is the presence of LIV in somatosensory cortex (S1), which is instead missing in M1. To analyze this, we have chosen a coronal section at -0.10 mm anterior-posterior (A.P.) from *bregma* and positioned the OF next to M1 or S1 (*Figure 4A*). Obtained *f* and *β* fields are shown in *Figure 4B* for two nominally identical slices from two different mice, while the *ρ* field is displayed in *Figure 4C*, with the iso-surfaces overlaid and their 3D configuration.

**Figure 4:**
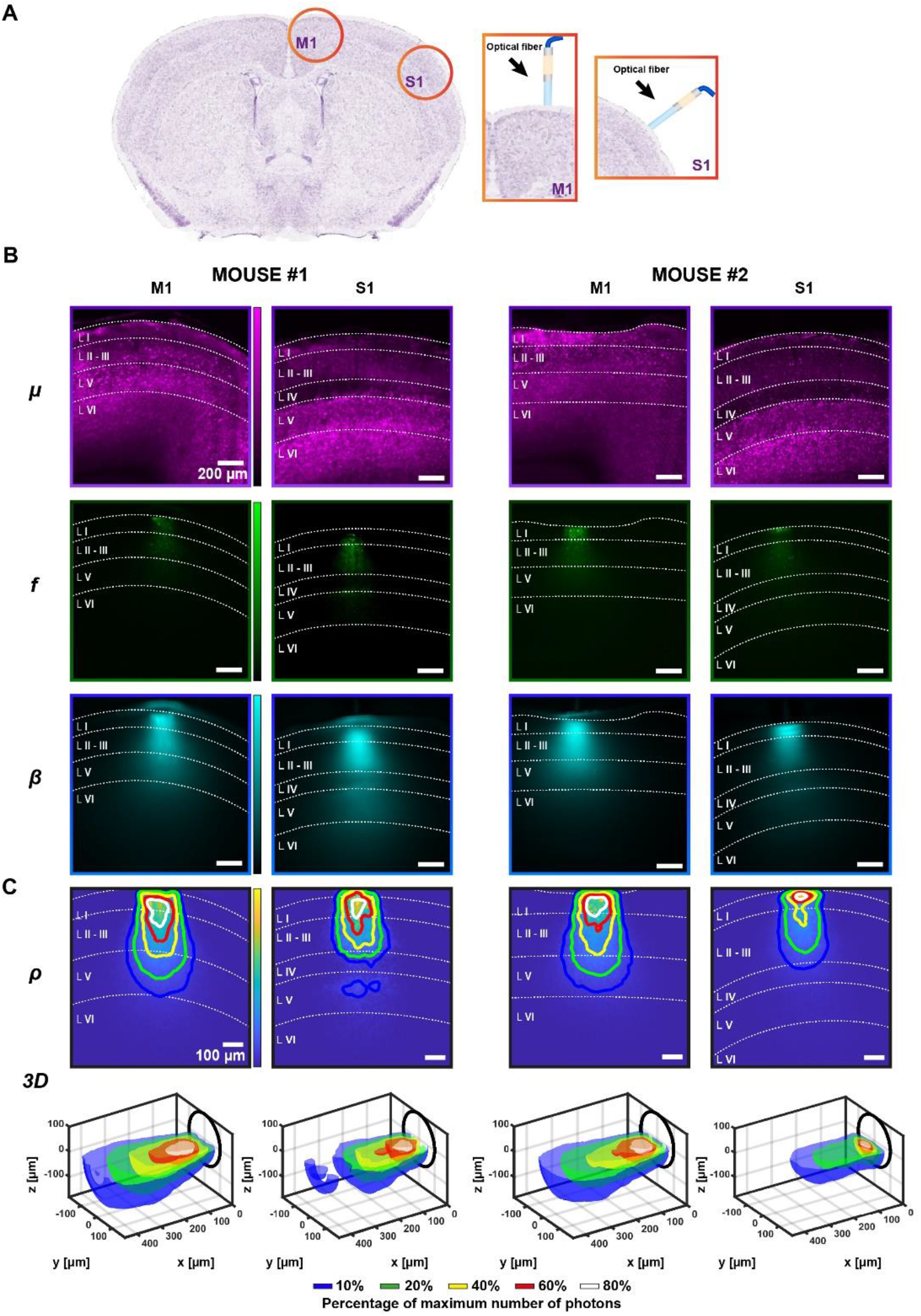
(**A**) Schematic representation of the optical fiber positioned next to M1 or S1. (**B**) Representative *µscope* PMT image (*top*), *fiber* PMT image (*middle*) and illumination field (*bottom*) of M1 and S1 for mouse#1 and mouse#2. Scalebar in all the panels (**B)** is 200 µm. (**C**) *(top)* Photometry collection efficiency field for each region with comparison of iso-intensity surfaces at 10%, 20%, 40%, 60%, and 80% of the maximum number of photons are shown (in blue, green, yellow, red, white respectively), *(bottom)* their 3D configuration as surfaces of revolution obtained by rotating the isolines around the fiber axis. Scalebar in all the panels (**C)** (*top*) is 100 µm. Images of *µ* and *f* in panel (**A**) were adjusted for visualization sake.

In this specific area of M1 the 10% intensity line reaches LV, which accounts for about the ∼11% of the overall collected signal (see bar graphs in *Figure 5A*), while for LI and LII-III we have found this value to be ∼30% and ∼60%, respectively. In S1, instead, the 10% isoline stops across the boundary between LII-III and LIV, resulting in a strong reduction (∼36%) of the influence of LV on the collected fluorescence, which accounts for the ∼7% of the overall fluorescence. The main contribution derives instead from LI and LII-III, which together generate more than the 80% of the photometry field signal. In *Thy1-* GCaMP6 transgenic line, LIV cell bodies do not express GCaMP6 [28], pyramidal neurons are smaller [30] and do not show a dendritic arborization toward LI and LII-III, with respect to typical LV pyramidal neurons [31]. Therefore, LIV acts as a shield for optical signal from deeper regions, and its thickness influences the ability to collect fluorescence from LV. This is shown by a comparison by 10% iso-intensity collection lines from S1 in mouse #2 and #1, with this latter showing some signal emerging from LV. A more detailed analysis of LIV thickness, displayed in *Figure 5B*, shows that LIV is slightly thinner (∼20 µm) in mouse #1, enabling more signal to reach the fiber facet from LV with respect to mouse #2.

**Figure 5:**
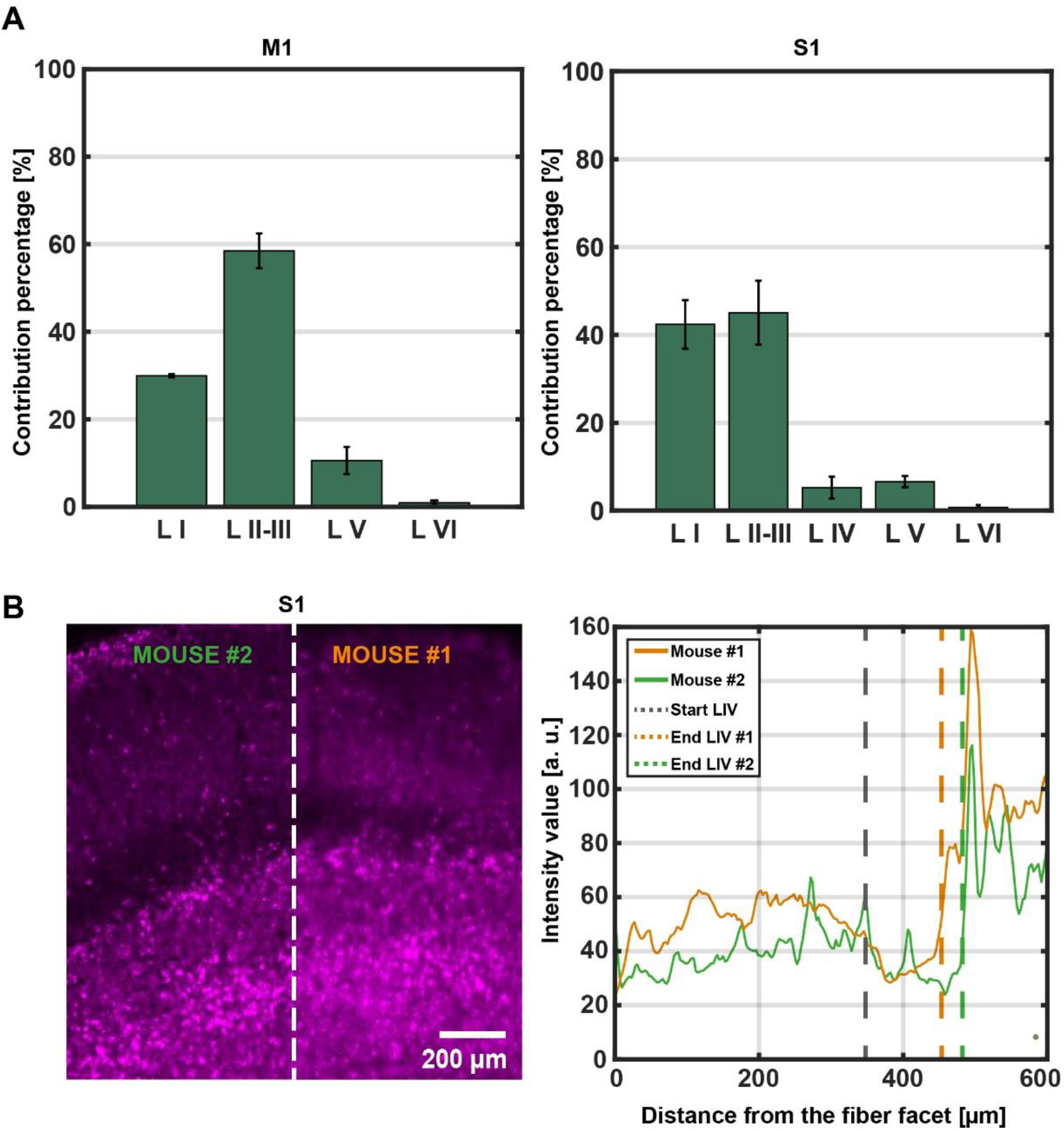
(**A**) Anatomical feature contributes for M1 and S1. (**B**) S1 microscopy images side-by-side with the thickness profile of Layer IV for mouse#1 with respect to mouse#2. Error bars in bar graphs in panels (**A**) represent the standard deviation of the percentages evaluated on N = 2 brain slices.

## DISCUSSION AND CONCLUSION

As fiber photometry is widely employed for collecting functional fluorescence from the mouse brain in free-moving animals, the definition of the collection volume was so far mainly based on (i) global properties of scattering in brain tissue, (ii) the distribution of the functional fluorescence in the targeted subpopulation of cells, and (iii) the collection properties of the employed device. We here report the evidence that an additional layer of complexity related to the specific anatomical features of the targeted brain structures and substructures has to be considered. The cytoarchitecture of different brain regions is crucial in defining the shape and the size of the light collection volume and, therefore, the type and number of cells contributing to the recorded signal. This is clearly shown in *Figure*

*3* for the transgenic mouse line *Thy1-* GCaMP6*s*, highlighting that the anatomy of the brain region under investigation directly influences the depth at which fluorescence is collected. Indeed, in M1, S1 and HP the thickness of the different layers set constrains or favors the ability to mine signal below a depth of 300 µm (*Figure 3A-C, Figure 4*). On the contrary, structures with more uniform cell distribution at the millimeters scale, like the striatum, show an even decrease of signal intensity as a function of depth (*Figure 3D*). Overall, this is the consequence of the heterogeneous optical properties of the brain, which are due to different cellular types, densities and shapes, and to local variations of the refractive index, which can also generate unexpected reflection and distortions when light travels across multiple regions [32]. On this respect, even small differences across multiple mice could result in detectable light collection differences and influence the experimental statistics, as observed in the comparison for S1 in two different animals (*Figure 4* and *5)* where a small difference in thickness of LIV affects light collection efficiency from LV. In this specific case, the effect is more pronounced due to the low expression of the *Thy1* promoter in LIV, and it highlights how the shape and the size of the collection volume is defined by the combination of: (i) the influence of the anatomy of the brain region of interest on the photometry field and (ii) the fluorescence distribution across the cell type of interest.

The method employed in this work can be extended to other transgenic mouse lines to identify the actual volumes contributing to the effective functional fluorescence, to better correlate recorded signals and their interpretation within the microcircuits of interest. Recent works have indeed shown how precise targeting of the region(s) and cell type of interests allows dissecting specific neural circuits related to memory, fear and epileptogenic activity, enabling to relate them to specific behavioral activity [33–38].

As well, the presented data highlight the limits of flat-cleaved optical fibers to collect photons below the first layers of cortex, and the need of developing complementary methods to achieve this aim. One example on this respect are tapered optical fibers [39], which have shown a more homogeneous signal distribution along a depth of a few millimeters and full compatibility with recently implemented photons detection methods like time-correlated single photon counting for lifetime fluorescence photometry [40]. This also highlights the need for novel technological paradigms for functional fluorescence collection in free-moving mice, able to better match the sensitivity volume with the anatomy of the brain structure of interest.

## MATERIALS AND METHODS

### Flat-cleaved optical fibers fabrication process

We realized flat-cleaved optical fibers stubs from 0.39 NA multimode optical fiber with core and cladding diameters of 200 μm and 225 μm, respectively (Thorlabs FT200UMT). Stubs were cutted with a manual fiber cleaver and connectorized to a 1.25 mm stainless-steel ferrule. The connectorized ends of the fiber stubs were manually polished. Patch fibers were realized from the same fiber type, using a SMA connector on one end and a 1.25 mm stainless-steel ferrule on the other end. Details of the procedure are provided in previous work [9].

### Optical setup specifications

The setup used to measure the illumination and the collection fields of an optical fiber is schematically shown in *Figure 1A*. A Pockels cell (Conoptics 350-80-02) is used to modulate the power of a λ_ex_ = 920 nm fs-pulsed near-infrared (NIR) laser beam (Coherent Chameleon Discovery). A quarter wave plate (Thorlabs AQWP05M-980) converts the linear polarization of the laser beam into circular polarization, and the beam diameter is 5-fold expanded and scanned in the xy plane by using a virtually conjugated galvo pair (Sutter). A 4X/0.28NA objective (Olympus XLFLUOR-340 4x/NA 0.28) is mounted on a z-axis fast piezo focuser (Phisik Instrument P-725.4CD). Fluorescence signal excited into coronal brain slices obtained from *Thy1*-GCaMP6*s* transgenic mice is re-collected by the same objective, routed on a non-descanned collection path through a dichroic mirror (Semrock FF665-Di02), two spherical lenses (Thorlabs LA1708-A and LA1805-A), and a bandpass filter (BPF, Semrock FF01-520/70-25), and detected by a photomultiplier tube (PMT, Hamamatsu H10770PA-40, the “μscope PMT”). The fiber stubs collecting fluorescent light were butt-coupled to a patch fiber and the light back-emitted from the patch fiber was collected by a microscope objective (Olympus Plan N 40x); a BPF (Semrock FF03-525/50-25) select the spectral region of interest and two spherical lenses (Thorlabs LA1050-A and LA1805-A) and a PMT (Hamamatsu H7422P-40, the “fiber PMT”), were used to measure the light intensity. Light emission diagrams at 473 nm (laser light from Laser Quantum Ciel) were imaged with a tube lens (Olympus U-TLU) and a sCMOS camera (Hamamatsu Orca Flash lite 4.0); light emission diagrams were registered over the light collection diagram by rescaling and roto-translation.

### Slices preparation

All experimental manipulations on mice were performed in accordance with protocols approved by Italian Ministry of Health. *Thy1*-GCaMP6*s* transgenic mice were anesthetized with isoflurane and were perfused transcardially with 4% paraformaldehyde (PFA) in 0.1 M sodium phosphate buffer. Brains were fixed for 24 h at 4 °C, washed in phosphate buffer saline (PBS) and sectioned (300 μm) coronally using a vibratome (Leica VT1000s). To perform the measurements in the hippocampus and in the striatum, the cerebral cortex and corpus callosum were removed manually with a razor blade.

### Data analysis

Data analysis was performed with custom written Matlab scripts. Source codes are available from the corresponding author on reasonable request.

Briefly, a background subtraction was performed on images acquired by the *fiber* PMT (*f*) and each slice was divided by the correspondent slice acquired by the *µscope* PMT (*µ*) to compensate for expression of fluorophore unevenness, obtaining *η*; the normalized average projection within the depth of focus volume was calculated and the so obtained field was multiplied by the normalized illumination image (*β*) to obtain *ρ*. Isosurfaces at 10%, 20%, 40%, 60% and 80% were evaluated on a smoothed (smooth window = 11) version of *ρ*.

Histograms reporting feature contributes to the collected signal were evaluated as percentage of the integral over the axial profile of the photometry efficiency fields within the anatomical structure (according to its depth). Error bars represent the standard deviation of the percentages evaluated on different brain slices.

## AUTHOR APPROVALS

All authors listed have made a substantial, direct and intellectual contribution to the work, and approved it for publication.

## COMPETING INTEREST STATEMENT

MDV, F. Pisanello are founders and hold private equity in Optogenix, a company that develops, produces and sells technologies to deliver light into the brain. Tapered fibers commercially available from Optogenix were used as tools in the research.

