## Supplementary Material for "Influence of anatomical features of different brain regions on the spatial localization of fiber photometry signals"

### Supplementary figures

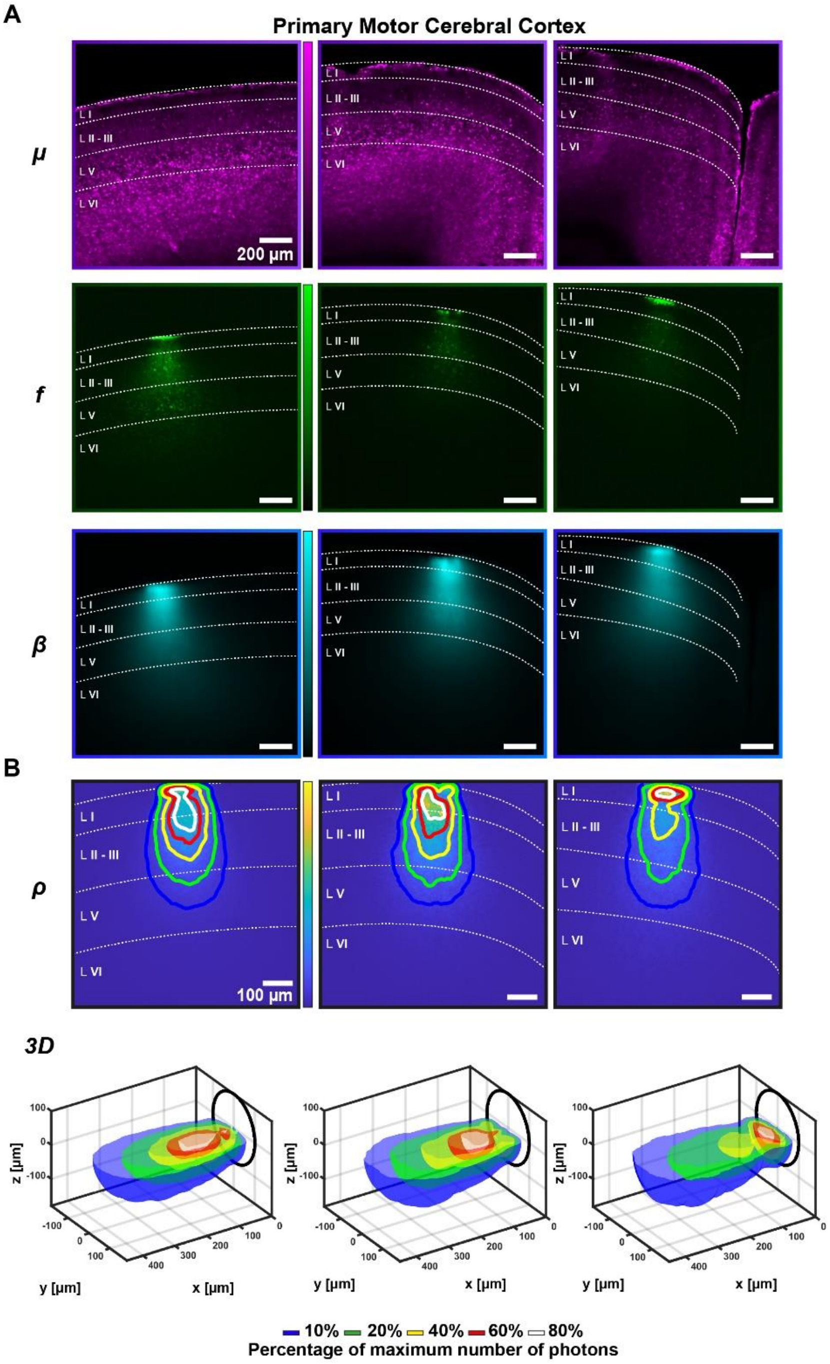


**Supplementary Figure 1: (A)** Representative *μscope* PMT image, *fiber* PMT image and the illumination field of primary motor cerebral cortex. Scalebar in all the panels **(A)** is 200 μm. **(B)** *(top)* Photometry collection efficiency field with comparison of iso-intensity surfaces at 10%, 20%, 40%, 60%, and 80% of the maximum number of photons are shown (in blue, green, yellow, red, white respectively); *(bottom)* their 3D configuration as surfaces of revolution obtained by rotating the isolines around the fiber axis. Scalebar in all the panels **(B)** (*top*) is 100 μm. Images of $\mu$ and $f$ in panel **(A)** were adjusted for visualization sake.

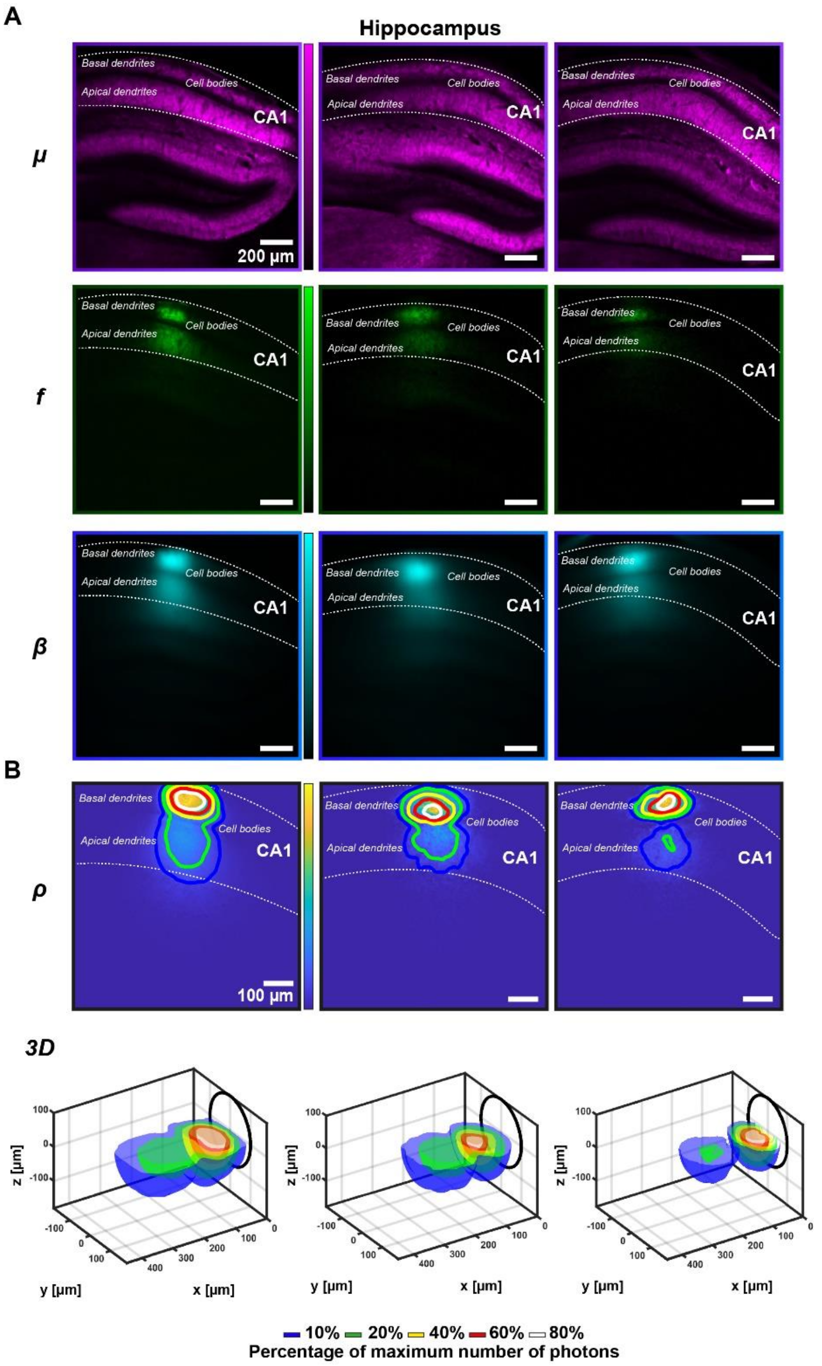


**Supplementary Figure 2: (A)** Representative *μscope* PMT image, *fiber* PMT image and the illumination field of hippocampus. Scalebar in all the panels **(A)** is 200 μm. **(B)** (*top*) Photometry collection efficiency field with comparison of iso-intensity surfaces at 10%, 20%, 40%, 60%, and 80% of the maximum number of photons are shown (in blue, green, yellow, red, white respectively); (*bottom*) their 3D configuration as surfaces of revolution obtained by rotating the isolines around the fiber axis. Scalebar in all the panels **(B)** (*top*) is 100 μm. Images of μ and f in panel **(A)** were adjusted for visualization sake.

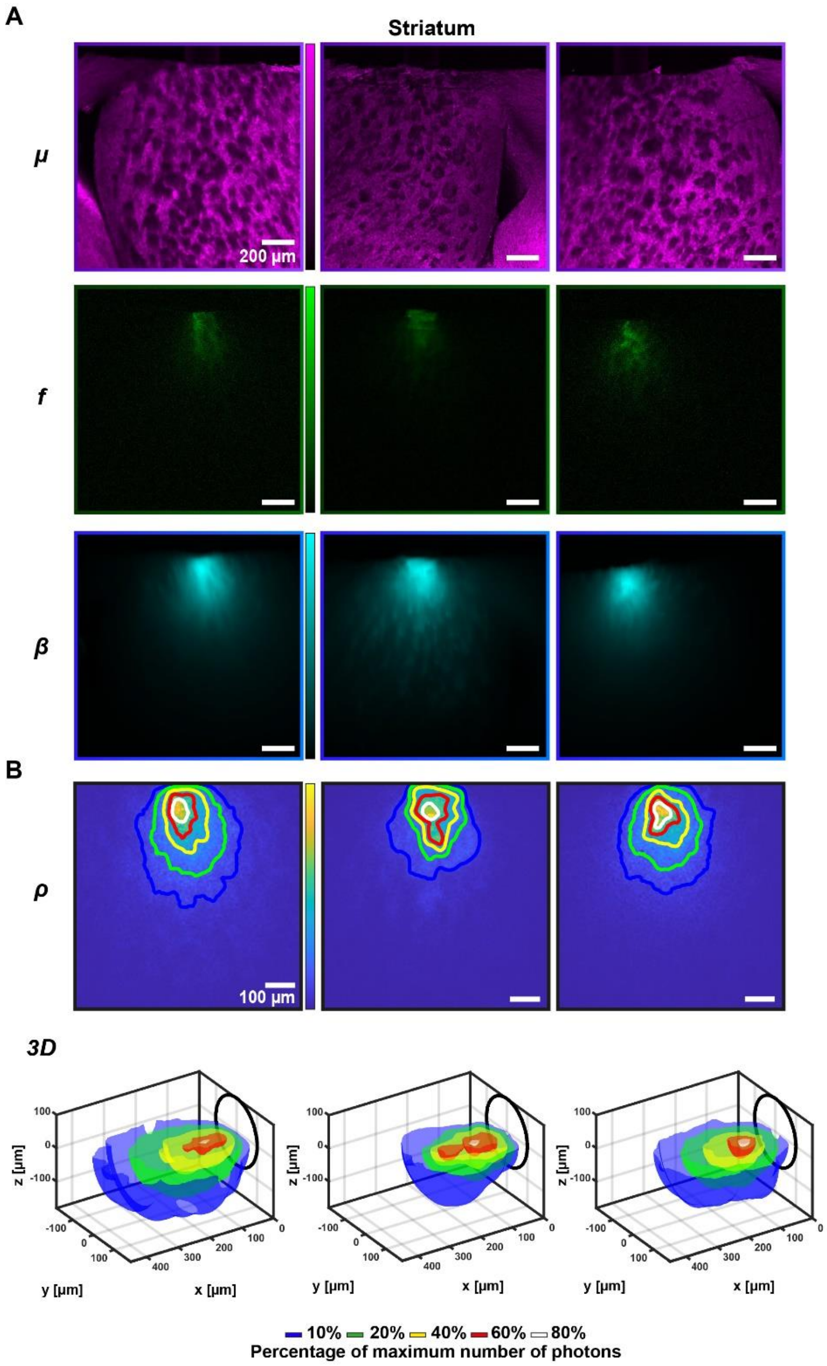


**Supplementary Figure 3: (A)** Representative *μscope* PMT image, *fiber* PMT image and the illumination field of striatum. Scalebar in all the panels **(A)** is 200 μm. **(B)** (*top*) Photometry collection efficiency field with comparison of iso-intensity surfaces at 10%, 20%, 40%, 60%, and 80% of the maximum number of photons are shown (in blue, green, yellow, red, white respectively); (*bottom*) their 3D configuration as surfaces of revolution obtained by rotating the isolines around the fiber axis. Scalebar in all the panels **(B)** (*top*) is 100 μm. Images of μ and f in panel **(A)** were adjusted for visualization sake.
